# Carbapenemases on the move: it’s good to be on ICE

**DOI:** 10.1101/392894

**Authors:** João Botelho, Adam P. Roberts, Ricardo León-Sampedro, Filipa Grosso, Luísa Peixe

## Abstract

The evolution and spread of antibiotic resistance is often mediated by mobile genetic elements. Integrative and conjugative elements (ICEs) are the most abundant conjugative elements among prokaryotes. However, the contribution of ICEs to horizontal gene transfer of antibiotic resistance has been largely unexplored. Here we report that ICEs belonging to mating-pair formation (MPF) classes G and T are highly prevalent among the opportunistic pathogen *Pseudomonas aeruginosa*, contributing to the spread of carbapenemase-encoding genes (CEGs). Most CEGs of the MPF_G_ class were encoded within class I integrons, which co-harbour genes conferring resistance to other antibiotics. The majority of the integrons were located within Tn*3*-like and composite transposons. A conserved attachment site could be predicted for the MPF_G_class ICEs. MPF_T_class ICEs carried the CEGs within composite transposons which were not associated with integrons. The data presented here provides aglobal snapshot of the different CEG-harbouring ICEs and sheds light on the underappreciated contribution of these elements for the evolution and dissemination of antibiotic resistance on *P. aeruginosa*.

## Introduction

Among the non-fermenting Gram-negative bacteria, the *Pseudomonas* genus is the one with the highest number of species [1, 2]. *Pseudomonas aeruginosa*, an opportunistic human pathogen associated with an ever-widening array of life-threatening acute and chronic infections, is the most clinically relevant species within this genus [3–5]. *P. aeruginosa* is one of the CDC “ESKAPE” pathogens – *Enterococcus faecium*, *Staphylococcus aureus*, *Klebsiellapneumoniae*, *Acinetobacter baumannii*, *P. aeruginosa* and *Enterobacter* species –,emphasizing its impact on hospital infections and the ability of this microorganism to “escape”the activity of antibacterial drugs [6]. *P. aeruginosa* can develop resistance to a wide range of antibiotics due to a combination of intrinsic, adaptive, and acquired resistance mechanisms,such as the reduction of its outer membrane permeability, over-expression of constitutive or inducible efflux pumps, overproduction of AmpC cephalosporinase, and the acquisition of antibiotic resistance genes (ARGs) through horizontal gene transfer (HGT) [4, 7, 8]. *P.aeruginosa* has a non-clonal population structure, punctuated by specific sequence types(STs) that are globally disseminated and frequently linked to the dissemination of ARGs [4, 9]. These STs have been designated as high-risk clones, of which major examples are ST111,61 ST175, ST235 and ST244.

Due to its high importance for human medicine, carbapenems are considered by the WorldHealth Organization (WHO) as Critically-Important Antimicrobials that should be reserved forthe treatment of human infections caused by MDR Gram-negative bacteria [10], such as *P.aeruginosa*. Carbapenem-resistant *P. aeruginosa* is in the “critical” category of the WHO’spriority list of bacterial pathogens for which research and development into new antibiotics is urgently required [11]. Besides *P. aeruginosa*, carbapenem resistance has been reported in other *Pseudomonas* spp. and is often mediated by the acquisition of carbapenemase-encoding genes (CEGs) [12–14]. Carbapenemases are able to hydrolyse carbapenems and confer resistance to virtually all ß-lactam antibiotics [15]. When it comes to the *Pseudomonas*genus, CEGs are mostly present on class I integrons within the chromosome [4]. Class Iintegrons are genetic elements that carry ARGs and an integrase gene, which controls integration and excision of genes [16–18]. Mobile genetic elements (MGEs) such as transposons, plasmids and integrative and conjugative elements (ICEs), are responsible forthe spread of ARGs [19–23]. The phage-inducible chromosomal islands are a recently reported family of MGEs, but unrelated to the carriage of ARGs [24].

Usually, the genes acquired by HGT are integrated in common hotspots in the host’schromosome, comprising a cluster of genes designated by genomic islands (GIs) [19, 25, 26]. This broad definition may also encompass other MGEs, such as ICEs and prophages. Although the exact origin of these elements remains unknown, a growing body of evidence shows that phages are one of the likely major ancestors of ICEs [27] [28]. ICEs are self-transmissible mosaic and modular MGEs that combine features of transposons and phages (ICEs can integrate into and excise from the chromosome), and plasmids (ICEs can also exist as circular extrachromosomal elements, replicate autonomously and be transferred by conjugation) [21,25, 29–31]. Integrative and mobilizable elements (IMEs) encode their own integration and excision systems, but take advantage of the conjugation machinery of co-resident conjugative elements to be successfully transferred [32]. ICEs usually replicate as part of the host genome and are vertically inherited, remaining quiescent, and with most mobility genes repressed [33,34]. These elements also encode recombinases related to those in phages and other transposable elements. Conjugation involves three mandatory components: a relaxase(MOB), a T4SS and a type-IV coupling protein (T4CP) [35, 36]. Four mating-pair formation(MPF) classes cover the T4SS among Proteobacteria: MPF_T_, MPF_G_, MPF_F_ and MPF_I_ [37]. The first is widely disseminated among conjugative plasmids and ICEs, while MPF_F_ is moreprevalent in plasmids of γ-Proteobacteria and MPF_G_ is found essentially on ICEs. MPF_I_ is rarelyidentified. Guglielmini *et al*. constructed a phylogenetic tree of VirB4, a highly conserved ATPase from the T4SS apparatus of different conjugative plasmids and ICEs, and formulated the hypothesis of interchangeable conjugation modules along their evolutionary history [38].A close interplay between these elements in the ancient clades of the phylogenetic tree was observed, suggesting that plasmids may behave like ICEs and vice-versa, reinforcing the common assumption that the line separating ICEs and conjugative plasmids is blurring [30,39]. These authors also searched more than 1000 genomes and found that ICEs are present in most bacterial clades and are more prevalent than conjugative plasmids [38]. It was also observed that the larger the genome, the higher the likelihood to harbour a conjugative element at a given moment, which supports the common assumption that bacteria with large genomes are more prone to acquire genes by HGT [40, 41].

Delimiting ICEs in genomic data remains particularly challenging [26]. Some signatures features are frequently observed, such as a sporadic distribution, sequence composition bias,insertion next to or within a tRNA gene, bordering attachment (*att*) sites and over-representation of mobility genes of the type-IV secretion system (T4SS). However, some ICEs present atypical features and may not be detected by these approaches [26, 40]. In *P.aeruginosa*, most ICEs fall into three large families: the ICE*clc*, pKLC102 and Tn*4371*. ThePAGI2(C), PAGI3(SG), PAGI-13, PAGI-15 and PAGI-16 were previously described as membersof the ICE*clc* family, while the PAPI-1, PAPI-2, PAGI-4 and PAGI-5 were linked to the pKLC102family [19]. The ICE_Tn*4371*_ family also represents a large group of ICEs with a common backbone and which are widely distributed, such as in *P. aeruginosa* UCBPP-PA14, PA7 and PACS171bstrains [21]. These ICEs have been frequently implicated in virulence [42, 43].

Previous reports characterized the complete nucleotide sequence of extra-chromosomal genetic elements housing different CEGs in pseudomonads [20, 44–47]; however, the association of CEGs with chromosome-located MGEs has rarely been investigated [48–50]. Taking into consideration that i) in pseudomonads, CEGs are frequently located within the chromosome, ii) ICEs are the most abundant conjugative elements in prokaryotes and iii) ICEs are more frequently identified in large bacterial genomes, such as in pseudomonads, we hypothesize that ICEs may play a key role in the horizontal spread of CEGs. To investigate this hypothesis, we developed an *in silico* approach to explore the association between ICEs and CEGs in pseudomonads.

## Methods

### Carbapenemases database

Antimicrobial resistance translated sequences were retrieved from the Bacterial Antimicrobial Resistance Reference Gene Database available on NCBI(ftp://ftp.ncbi.nlm.nih.gov/pathogen/Antimicrobial_resistance/AMRFinder/data/2018-04-16.1/). The resulting 4250 proteins were narrowed down to 695 different carbapenemases to create a binary DIAMOND (v. 0.9.21, https://github.com/bbuchfink/diamond) database [51]. Only the sequences presenting ‘carbapenem-hydrolyzing’ or ‘metallo-beta-lactamase’ on fasta-headers were used to build this local database.

### Genome collection and blast search

A total of 4565 *Pseudomonas* genomes was downloaded from NCBI (accessed on the 24^th^ ofApril, 2018). These genomes were blasted against the local carbapenemase database using the following command: ‘diamond blastx –d DB.dmnd –o hits.txt --id 100 --subject-cover 100-f 6 --sensitive’.

### Bioinformatic prediction of ICEs and genetic environment analyses

The CEG-harbouring *Pseudomonas* genomes were annotated through Prokka v. 1.12(https://github.com/tseemann/prokka) [52]. The translated coding sequences were analysed in TXSScan/CONJscan platform to inspect the presence of ICEs(https://galaxy.pasteur.fr/root?tool_id=toolshed.pasteur.fr%2Frepos%2Fodoppelt%2Fconjscan%2FConjScan%2F1.0.2) [37]. All ICEs harbouring CEGs predicted by TXSScan/CONJscan were inspected for direct repeats that define the boundaries of the element. The complete nucleotide sequence in Genbank format of corresponding records was imported into Geneious v. 9.1.8 to help delimiting genomic regions flanking the ICEs [53]. Complete ICE sequences were aligned with EasyFig v. 2.2.2 (http://mjsull.github.io/Easyfig/files.html) [54]. Screening of complete ICEs for ARGs was achieved by ABRicate v. 0.8(https://github.com/tseemann/abricate). Phage and insertion sequences were inspectedthrough PHASTER (http://phaster.ca/) and ISfinder (https://www-is.biotoul.fr/), respectively[55, 56]. Multiple Antibiotic Resistance Annotator (MARA, http://mara.spokade.com) wasused to explore the genetic background of the CEGs [57]. Orthologous assignment and functional annotation of integrase sequences was achieved through EggNOG v. 4.5.1(http://eggnogdb.embl.de/#/app/home) and InterProScan 5(https://www.ebi.ac.uk/interpro/search/sequence-search) [58, 59].

### Phylogenomics

All CEG-harbouring *P. aeruginosa* genomes were mapped against the *P. aeruginosa* PAO1reference strain (accession number NC_002516.2), to infer a phylogeny based on the concatenated alignment of high quality single nucleotide polymorphisms (SNP) using CSIPhylogeny and standard settings [60]. The phylogenetic tree was plotted using the iTOLplatform (https://itol.embl.de/).

### MLST and taxonomic assignment of unidentified species

To predict the sequence type (ST) of the strains harbouring ICEs, the *P. aeruginosa* MLSTwebsite (https://pubmlst.org/paeruginosa/) developed by Keith Jolley and hosted at the University of Oxford was used [61]. Taxonomic assignment of unidentified species carrying ICEs was achieved by JSpeciesWS v. 3.0.17 (http://jspecies.ribohost.com/jspeciesws/#home)174 [62].

## Results

### A plethora of carbapenemase-encoding genes was identified in a subset of *Pseudomonas*species

From the total *Pseudomonas* genomes analysed (n=4565), 313 CEGs were identified in 297genomes (**Figure 1** and **Table S1**). As expected, *bla*_VIM-2_ represents the majority of the CEGs found among *Pseudomonas* spp., being detected mainly in *P. aeruginosa*, followed by *P.plecoglocissida*, *P. guariconensis*, *P. putida*, *P. stutzeri* and 16 genomes corresponding to unidentified species (**Table S1**). Curiously, some strains presented two CEGs, either presentinga duplication of the same gene, such as *bla*_IMP-34_ from NCGM 1900 and NCGM 1984 Japanese isolates, or harbouring different CEGs, such as *bla*_IMP-1_ and *bla*_DIM-1_ in isolates 97, 130 and 142recovered in Ghana (**Table S1**, highlighted in red). A wide variety of STs was also observed,including the high-risk clones ST111, ST175 and ST244.

**Figure 1.**
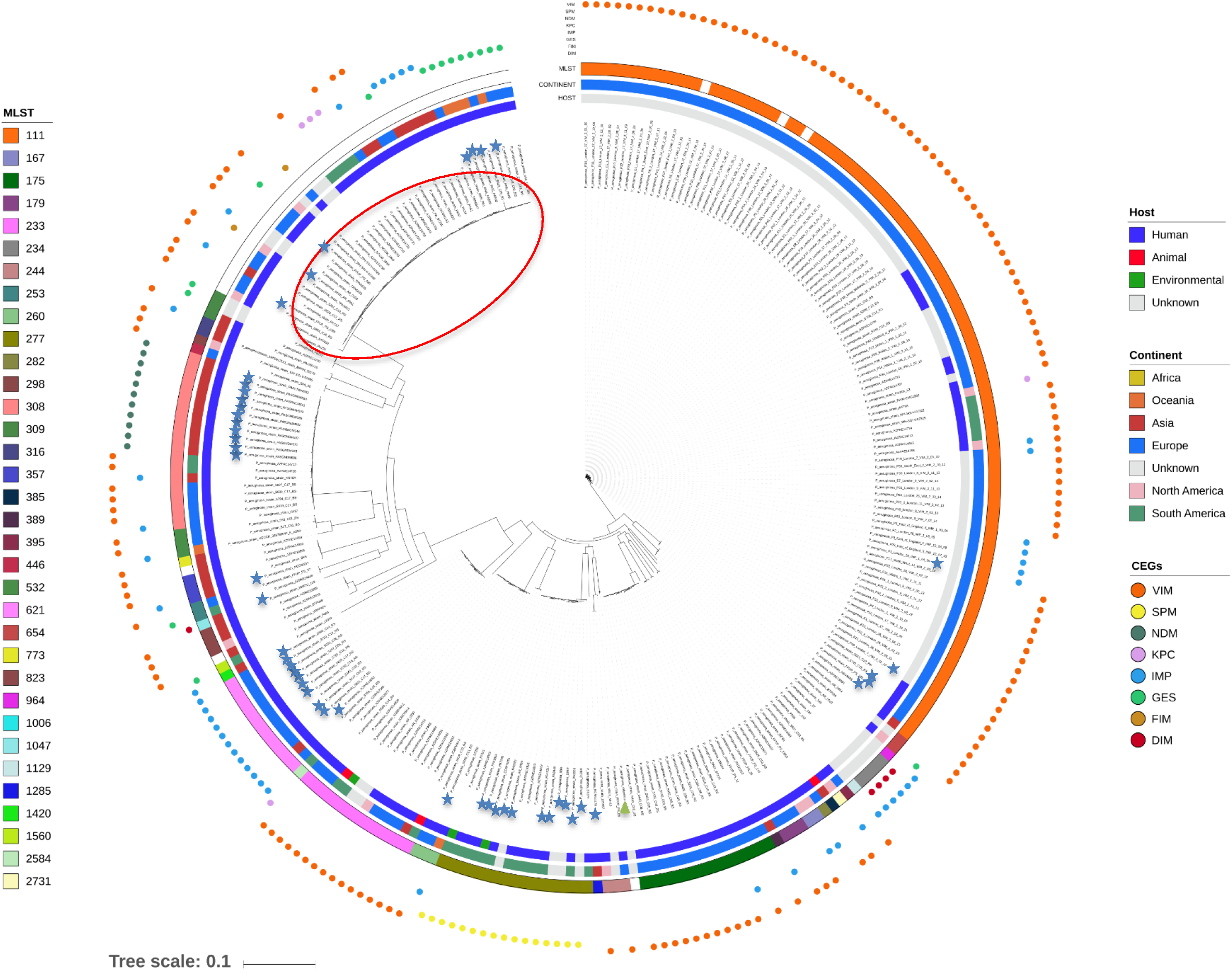
Whole-genome phylogeny of the CEG-carrying *P. aeruginosa* isolates. The maximum-likelihood phylogenetic tree was constructed using 146,106 SNPs spanning the whole genome and using the *P. aeruginosa* PAO1 genome (highlighted by a green triangle) asa reference. Multilocus sequence typing (MLST), continent and host data are reported on the outer-most, middle and inner-most circles, respectively. The strains belonging to a double ST profile (ST235/ST2613) are included within the red ellipse. Blue stars point out *P. aeruginosa*strains for which a CEG-harbouring ICE was predicted. The *P. aeruginosa* AR_0356 genome(accession number CP027169.1) was removed from the tree since it corresponds to a strain of which host and origin are unknown. The phylogenetic distance from the tree root to this genome is 1 (calculated with the tree scale). The Newick format file for the original tree is included in the Supplementary information.

### Detection of ICEs encoding carbapenemases in *Pseudomonas* spp

65.5% (205/313, **Table S1**) of the CEG hits are located within small contigs, with a sequence smaller than 20kb in length. The presence of repeated regions, such as those encoding for transposases, tend to split the genome when second-generation sequencing approaches are used. Based on information retrieved from NCBI (accessed on the 24^th^ of May, 2018), the total number of bacterial genomes sequenced at the chromosome/complete genome level is12,077, while the number of genomes sequenced at the scaffold/contig is much larger(127,231). With this sequencing limitation, we were still able to identify 49 ICEs associated with CEGs (n=20 with complete sequence) among all pseudomonads genomes (**Table 1**, **TableS1 and Figure 1**). When an ICE location was attributed to a CEG located on a small contig, the assumption was based on previously published data, as pointed out on **Table 1**. Besides theaforementioned ICEs, we also identified a putative MGE within *Pseudomonas* sp. NBRC111143 strain (**Table S1**). The T4CP-encoding gene was absent from this *bla*_IMP-10_-carryingelement, which could be due to contig fragmentation or gene absence. In case the gene isactually missing, this element could still be mobilized by the conjugation machinery of an ICEor conjugative plasmid(s) present in the host, and should be classified as an IME.

The ICEs identified here were all integrated within *P. aeruginosa* genomes (with the exceptionof the one element identified in *Pseudomonas* sp. PONIH3 genome) and AT-rich whencompared to their host’s chromosome; the mean GC value for this species is 66.2% accordingto EZBioCloud (https://www.ezbiocloud.net/taxon?tn=Pseudomonas%20aeruginosa) (**Table 1**).

**Table 1.**
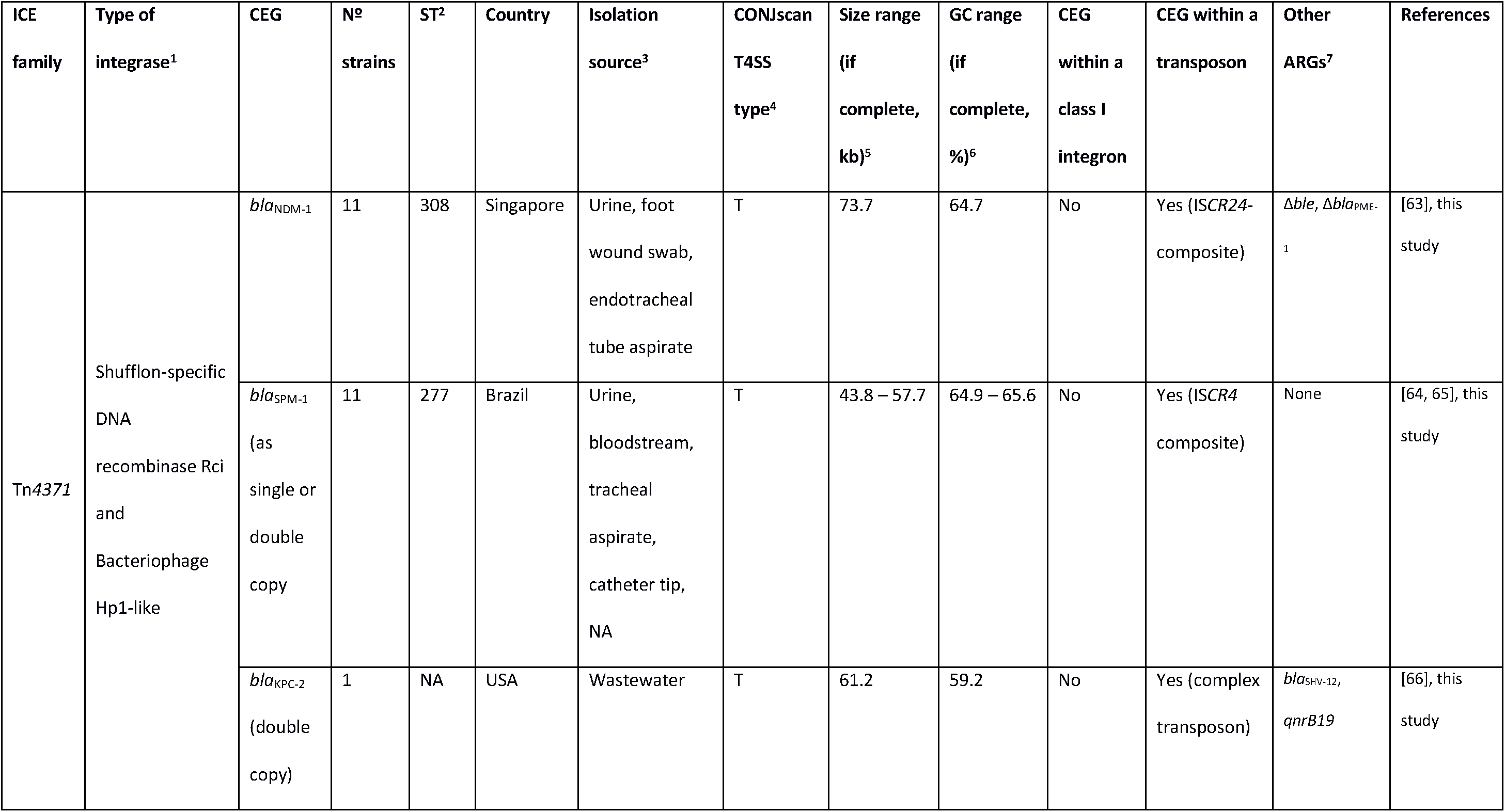
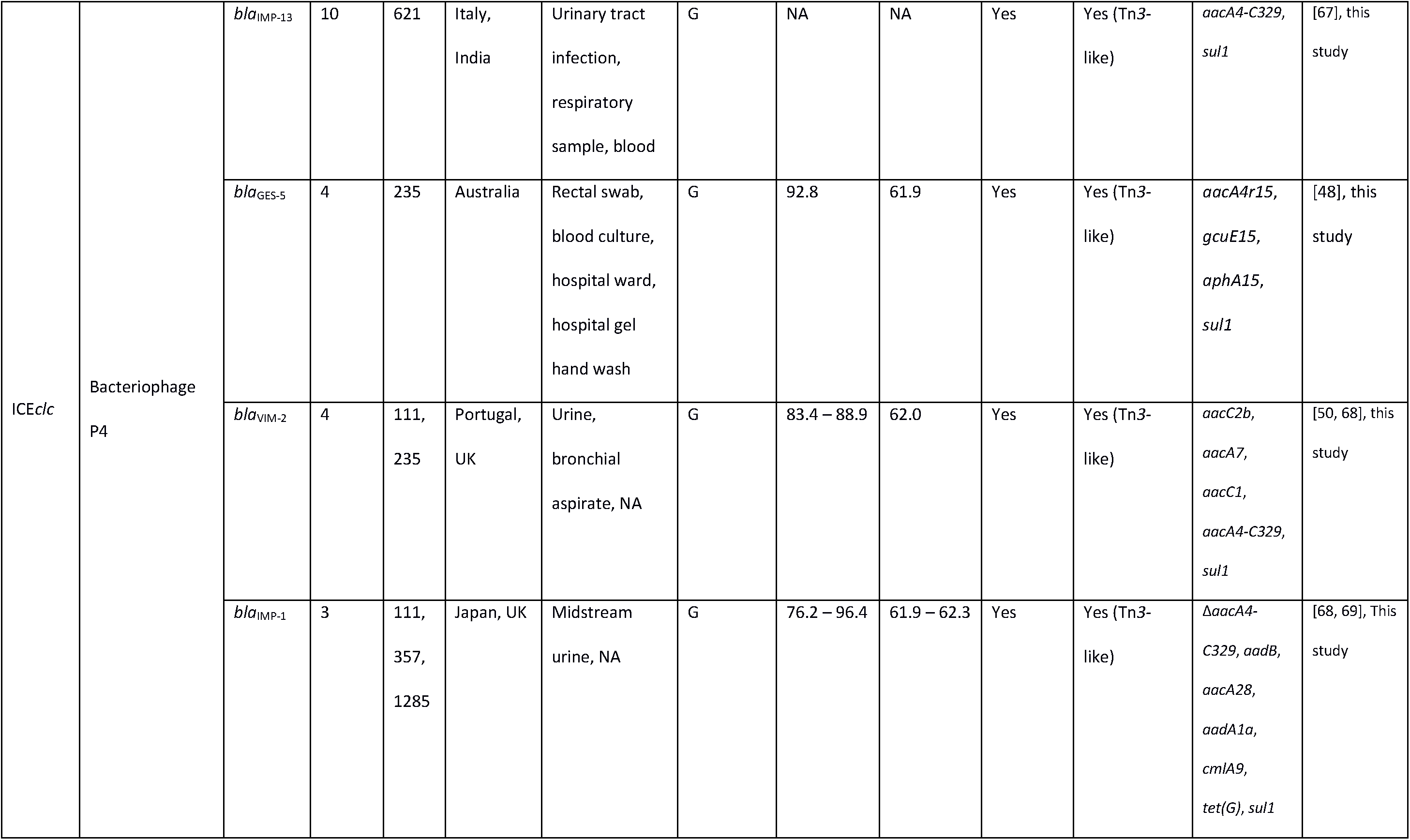
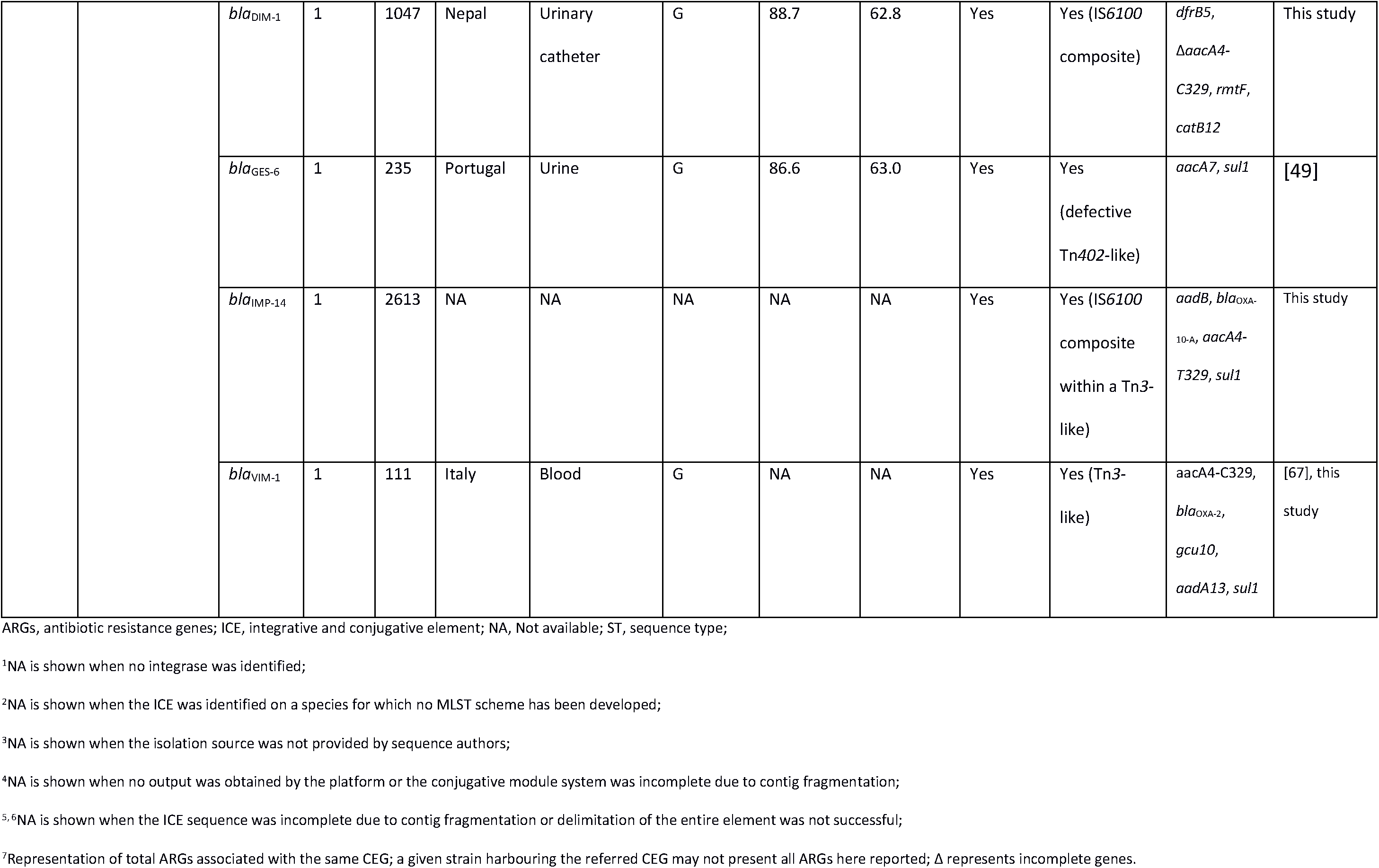
Main characteristics of CEG-carrying ICEs described in this study.

All ICEs identified here possessed only one tyrosine integrase (**Figure 2**). ICEs belonging to the ICE*clc* family (MPF_G_ class) carried an integrase belonging to the bacteriophage P4-like family, while ICEs belonging to the ICE_Tn*4371*_ family (MPF_T_ class) carried an integrase belonging to shufflon-specific DNA recombinase Rci and Bacteriophage Hp1-like family (**Table 1**).^31^ Rci and Hp1-like were only distantly related (13% amino acid identity) to P4-like integrases. Orthologous assignment of these integrases revealed that the former and the later integrases identified were present in more than 100 and 400 proteobacteria species, respectively. While P4-like integrases were more prevalent on γ-proteobacteria, half of the strains carrying Rci and Hp1-like integrases belong to the α-proteobacteria.

**Figure 2.**
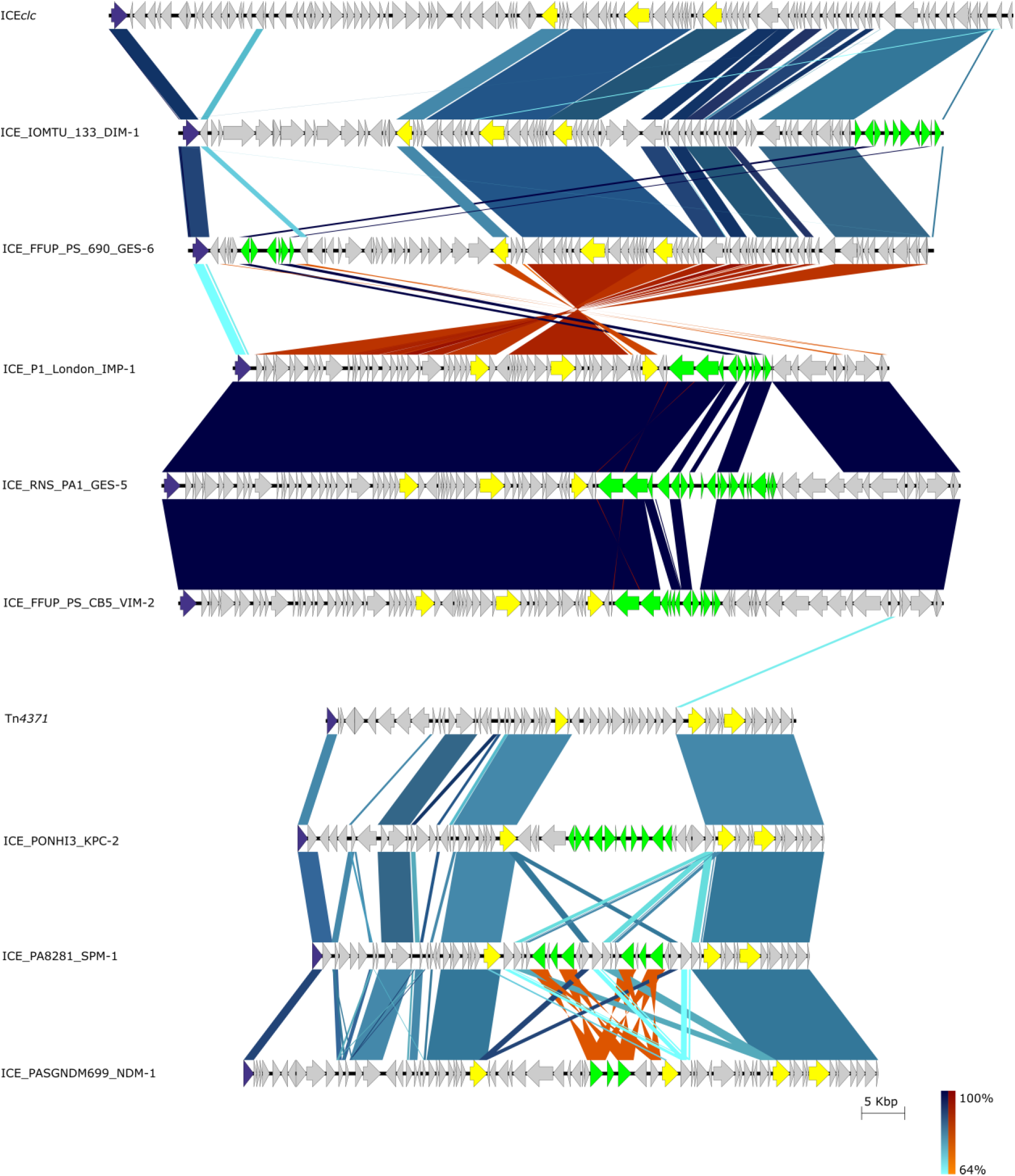
Blastn comparison among multiple ICEs described in this study. A gradient of blue and red colours is observed for normal and inverted BLAST matches, respectively. Model elements (ICE*clc* for the MPF^G^ and Tn*4371* for the MPF^T^ classes, respectively) were also included for comparison. The arrows and arrowheads point the orientation of the translated coding sequences. In purple are highlighted the integrases, in yellow the mandatory features of a conjugative system according to Cury *et al*. [40] and in green the transposons harbouring the CEGs. A more detailed view of some of these transposons is displayed in **Figure 3**.

We observed that MPF_G_ class ICEs tend to integrate next to a single copy of tRNA^G^^ly^ or a cluster of two tRNA^Glu^ and one tRNA^Gly^ genes, which is in agreement with previous findings [26, 40]. A conserved 8-bp *att* site (5′-CCGCTCCA) flanked all complete ICEs of the MPF_G_ class identified here (**Table 1**). Notably, most ICEs of this class were adjacent to phages (either at the 5’-or the 3’-end) targeting the same *att* site as the neighbour ICE. No *att* site could be identified for the integration of MPF_T_ class ICEs. A gene encoding for a catechol 1,2-dioxygenase and a gene encoding for a protein with no described conserved domain were found flanking the *bla*_SPM-1_-harbouring ICEs. Regarding the elements carrying *bla*_NDM-1_, a gene encoding for a different protein also with no conserved domain identified and a gene encoding for the type III secretion system adenylate cyclase effector ExoY were separated upon insertion of these ICEs. Integration next to hypothetical proteins or tRNA genes was commonly observed.

### Carbapenemases are frequently encoded within transposons

CEGs were associated with class I integrons frequently co-harbouring aminoglycoside resistance genes when associated with MPF class G ICEs (**Table 1**). Class I integrons were often associated with a wide array of transposons, such as the Tn*3* superfamily transposons and the IS*6100* composite elements (**Table 1**). MPF_T_ class ICEs were targeted by more complex elements, such as the composite transposons carrying *bla*_SPM-1_ and *bla*_NDM-1_ (**Table 1**). The *bla*_NDM-1_ gene was identified in Singapore in ICE_Tn*4371*_6385 and associated with ST308, as recently reported [63]. The *bla*_NDM-1_ was flanked by two IS*CR24*-like transposases. *bla*_SPM-1_ waslinked to ICE_Tn*4371*_6061, a recently described ICE [64]. Again, the CEG was located within an IS*CR4*-like composite transposon. IS*CR* elements are atypical elements of the IS*91* family which represent a well-recognized system of gene capture and mobilization by a rolling-circle transposition process [21, 70].

Besides previously described *bla*_NDM-1_ and *bla*_SPM-1_ harbouring ICEs, we characterize here new ICE elements of MPF_G_ and MPF_T_ classes (**Table 1** and **Figure 3**). The *bla*_DIM-1_-harbouring ICE from IOMTU 133 strain was integrated between the 3’-end of a tRNA^Gly^ gene (IOMTU133_RS11660) and a gene encoding for the R body protein RebB (IOMTU133_RS12085). *bla*_DIM-1_ was first described as a single gene cassette located within a class I integron associated with a 70-kb *Pseudomonas stutzeri* plasmid recovered in the Netherlands [13]. However, the integron carrying *bla*_DIM-1_ in strain IOMTU 133 was unrelated to the one from the *P. stutzeri* plasmid, harbouring genes encoding for aminoglycoside (*aacA4-C329* and *rmtf*), trimethoprim (*dfrB5*) and chloramphenicol (*catB12*) resistance (**Figure 3A**). Direct repeats (DRs) were found flanking the entire IS*6100* composite transposon (5’-TTCGAGTC), indicating the transposition of this element into the ICE element. Besides being identified as a composite transposon, IS*6100* was frequently observed as a single copy at the 3’end of the class I integron (**Figures 3B and 3C**), suggesting that these elements were derived from the In*4* lineage [71]. The *bla*_IMP-1_ from the NCGM257 strain identified in Japan belonged to a different ST (ST357) than the frequently identified ST235 associated with the spread of this CEG in this country [72]. The CEG was also shown to be associated with a novel complex class I integron, co-harbouring *aadB*, *cmlA9* and *tet(G)* genes encoding resistance to aminoglycosides, chloramphenicol and tetracyclines, respectively (**Figure 3B**). This integron was inserted (DRs 5’-GAGTC) within a mercury resistance transposon. This genetic organization was frequently recovered among other ICE-harbouring strains, such as the onesassociated with *bla*_GES-5_, *bla*_IMP-13_ and *bla*_IMP-14_ (**Table 1**). The entire ICE was integrated into the chromosome of NCGM257 strain between the 3’-end of a tRNA^Gly^ gene (PA257_RS24790) and the aforementioned *Pseudomonas* phage Pf1-like element. The new ICE identified on the P1_London_28_IMP_1_04_05 strain presented *bla*_IMP-1_ in a different In*4*-like integron than that observed for the NCGM257 strain, even though both elements were associated with a Tn*3*-like transposon (**Figure 3C**). Unlike most ICEs of the MPF_G_ class, its integration occurred between a gene encoding for a LysR family transcriptional regulator (AFJ02_RS19410) and a gene encoding for a hypothetical protein (AFJ02_RS19770). Regarding the *bla*_KPC-2_-harbouring *Pseudomonas* sp. PONHI3 strain, a tetra correlation search revealed that this strain was highly similar (Z-score above the 0.999 cut-off) to *Pseudomonas mosselii* SJ10 (accession number NZ_CP009365.1). Average nucleotide identity based on BLAST (ANIb) analysis of these genomes revealed that both strains belong to the same species, since the ANIb value was above the 95% cut-off for species delineation [73]. However, the ANIb value for both strains was below the cut-off when compared with the *P. mosselii* DSM 17497 type train (accession number NZ_JHYW00000000.1), suggesting that both strains may comprise novel species within the *Pseudomonas putida* phylogenetic group [2]. The PONHI3 strain carried a double copy of *bla*_KPC-2_ within an ICE from MPF_T_ class. A complex genetic environment was found surrounding these genes (**Figure 3D**). This ICE was integrated between a gene encoding for a biopolymer transport protein ExbD/TolR (C3F42_RS18665) and a gene encoding for an alpha/beta hydrolase (C3F42_RS18995).

**Figure 3.**
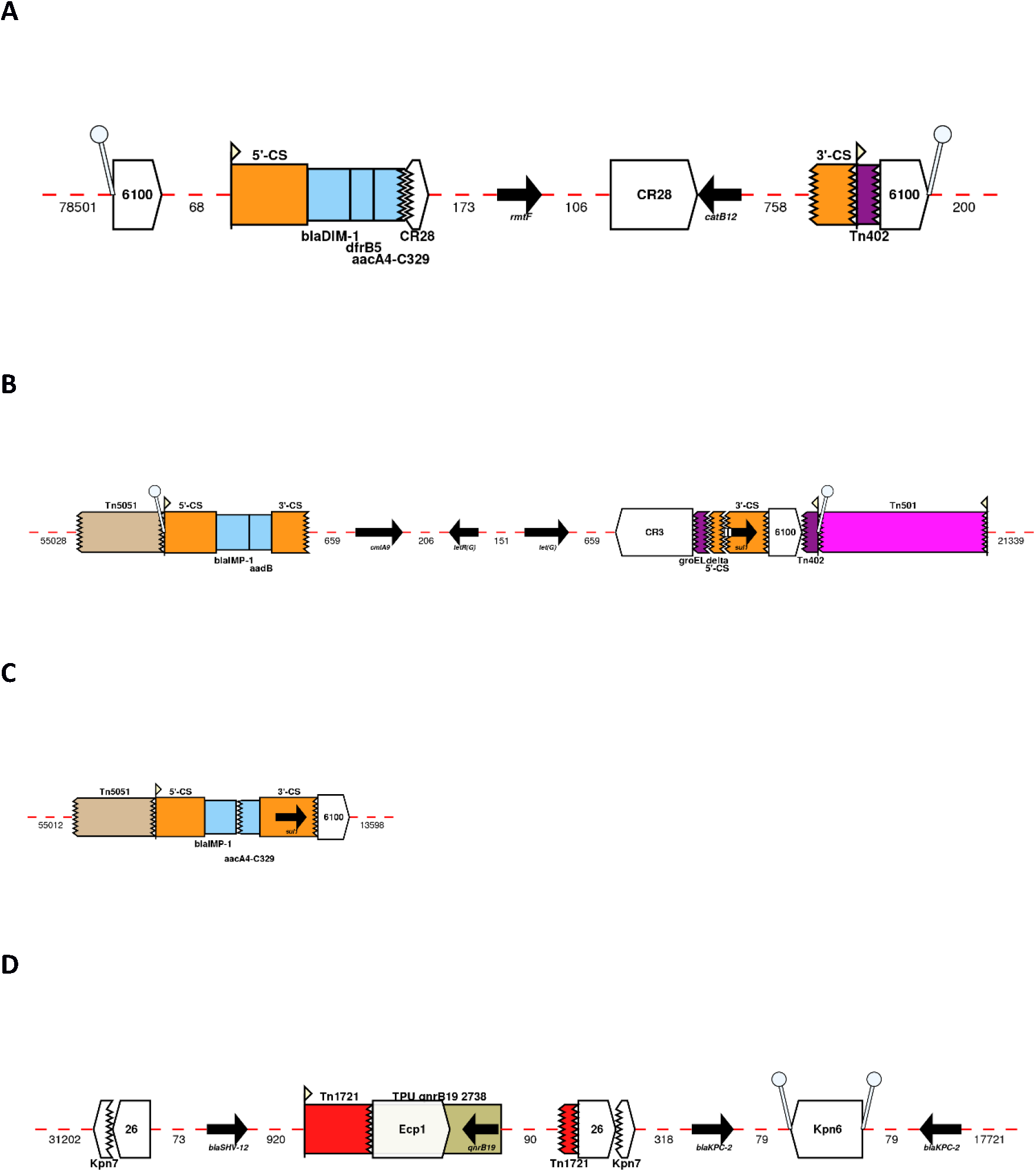
Genetic environment of novel ICEs harbouring *bla*_DIM-1_ (A), *bla*_IMP-1_ (B and C) and a double copy of *bla*_KPC-2_ (D). Arrows indicate the direction of transcription for genes. The dashed part of the arrow indicates which end is missing, for other features the missing end is shown by a zig-zag line. Gene cassettes are shown by pale blue boxes, the conserved sequences (5’ and 3’-CS) of integrons as orange boxes and insertion sequences as white block arrows labelled with the IS number/name, with the pointed end indicating the inverted right repeat (IRR). Gaps >50 bp are indicated by dashed red lines and the length in bp given. Unit transposons are shown as boxes of different colors and their IRs are shown as flags, with the flat side at the outer boundary of the transposon. Direct repeats are shown as ‘lollipops’ of the same color.

### An atypical GI encoding carbapenemases

Besides ICEs, we also identified an atypical 19.8-kb long GI harbouring *bla*_VIM-2_ in *P. aeruginosa* AZPAE13853 and AZPAE13858 strains from India (**Figure S1**). A similar element was also observed in *P. aeruginosa* BTP038 strain from the USA, with the exception that the Tn*402*-like transposon harbouring *bla*_VIM-2_ was orientated in an inverted position. Five base-pair DRs (5’-CTCTG in AZPAE13853 and AZPAE13858 and 5’-CTGAG in BTP038 strains) were found flanking this transposon structure. Importantly, in these strains the GIs were flanked by identical signal recognition particle RNAs (srpRNAs), indicating a strong site preference for these elements.

## Discussion

Our results show that *bla*_VIM_ and *bla*_IMP_ are widely disseminated, both geographically and phylogenetically (across *Pseudomonas* spp.). Moreover, and as previously described, *bla*_VIM-2_ was the most frequently reported CEG (**Figure 1** and **Table S1**) [4]. On the other hand, *bla*_SPM-1_ is still restricted to *P. aeruginosa* and Brazil (or patients who had been previously hospitalized in Brazil) [65]. Even though ST235 has been frequently linked to the dissemination of ARGs, no CEG-harbouring strains belonging to this ST were identified. Curiously, some strains (highlighted on **Figure 1**) belong to a double ST profile(ST235/ST2613), since the strains carry a double copy with different allele sequences of the house-keeping gene *acsA*, encoding for an acetyl-coenzyme A synthetase. These genes only display 80,3% nucleotide identity. We plan to conduct comparative genomic studies to explore the idiosyncrasies of these double ST profile strains.

Not all CEGs are likely to be geographically and phylogenetically disseminated, but those that are more promiscuous present a serious threat. The geographical distribution of the high-risk clones and the diversity of CEGs propose that the spread of these STs is global and the acquisition of the resistance genes is mainly local [4, 68]. Previous studies suggest that environmental species may pose an important reservoir for the dissemination of clinically relevant carbapenemases, which are vertically amplified upon transfer to *P. aeruginosa* high-risk clones [12, 14]. The high prevalence of these elements among high-risk clones may be partially explained by the genetic capitalism theory, given that a widely disseminated ST should have a greater probability of acquiring new CEGs and to be further selected and amplified due to the high antibiotic pressure in the hospital environment [74]. Other theories support that the high-risk clones have a naturally increased ability to acquire foreign DNA, since these STs appear to have lost the CRISPR (clustered regularly interspaced short palindromic repeats)-Cas (CRISPR associated proteins) system, which act as an adaptive immune system in prokaryotic cells and protects them from invasion by bacteriophages and plasmids [75–77].

This study underestimates the extent of host range because only ICEs in sequenced genomes were detected. Also, identification of new ICEs could only be achieved in complete genomes or contigs with a sequence length large enough to include the full (nor near complete) sequence of the ICE. As so, it is important to highlight the need to perform third generation sequencing on CEG-harbouring genomes to avoid fragmentation of the genetic environment surrounding the gene and to provide a wider view of complete supporting ICEs and other MGEs. All ICE elements here identified fulfilled the criteria to be considered conjugative as proposed by Cury *et al*.: a relaxase, a VirB4/TraU, a T4CP and minimum set of MPF type-specific genes [40]. ICEs tend to integrate within the host’s chromosome by the action of a tyrosine recombinase, even though some elements may use serine or DDE recombinases instead [30]. Though rare, some elements encode for more than one integrase, most likely resulting from independent integration of different MGEs [40]. Conserved sites are hotspots for ICE integration due to their high conservation among closely related bacteria, and so expanding the host range and be stably maintained after conjugative transfer [78, 79]. ICEs were often integrated next to phages highly similar to the *Pseudomonas* phage Pf1 (NC_001331.1), a class II filamentous bacteriophage belonging to the *Inoviridae* family [77]. Pf1-like phages are widely disseminated among *P. aeruginosa* strains and may have a role in bacterial evolution and virulence [80–82]. Interestingly, no representative of the pKLC102 family was linked to the dissemination of CEGs. This may be explained due to a higher affinity of the transposons carrying the CEGs for hotspots located within representatives of the other two families.

MGEs specifically targeting conserved regions of the genome such as tRNAs are common and this specificity represents an evolutionary strategy whereby the target site of an element is almost guaranteed to be present, due to its essentiality, and very unlikely to change due to biochemical constraints of the gene product. We think a similar situation exists for the elements found between the small srpRNAs described on the atypical GI element here identified and is in contrast to the more permissive nature of target site selection shown for example, by elements of the Tn*916*/Tn*1545* family [83].

Here, we revealed that different Tn*3*-like and composite transposons harbouring a wide array of CEGs were transposed into MPF G and T ICE classes, which were most likely responsible for the dissemination of these genes through HGT and/or clonal expansion of successful *Pseudomonas* clones. This study sheds light on the underappreciated contribution of ICEs for the spread of CEGs among pseudomonads (and potentially further afield). With the ever-growing number of third-generation sequenced genomes and the development of more sophisticated bioinformatics, the real contribution of these ICEs will likely rapidly emerge.

Recently, it was shown that interfering with the transposase-DNA complex architecture of a Tn*916*-like conjugative transposon (also known as ICE) lead to transposition inhibition to a new host [84]. In the future, it would be interesting to determine if the same mechanism is observed for tyrosine recombinases present in ICE*clc* and Tn*4371* derivatives, as well as in other MPF ICE classes, as a potential approach to interfere with the spread of antimicrobial resistance.

## Acknowledgments

This study received financial support from the European Union (FEDER funds POCI/01/0145/FEDER/007728) and National Funds (FCT/MEC, Fundação para a Ciência e Tecnologia and Ministério da Educação e Ciência) under the Partnership Agreement PT2020 UID/MULTI/04378/2013. J. B. and F. G. were supported by grants from Fundação para a Ciência e a Tecnologia (SFRH/BD/104095/2014 and SFRH/BPD/95556/2013, respectively). We thank Álvaro San Millan for helpful discussions. We also thank Benjamin Buchfink (DIAMOND), Jean Cury (TXSScan/CONJscan) and Sally Partridge (MARA) for their valuable assistance.

## Author contributions

JB, APR, FG and LP designed the study; JB and RLS performed the *in silico* analysis; JB wrote the manuscript. All the authors approved the final manuscript.

## Competing interests

None to declare.

